# Vision plays a calibrating role in discriminating threat-related vocal emotions

**DOI:** 10.1101/2023.09.27.559716

**Authors:** Federica Falagiarda, Valeria Occelli, Olivier Collignon

## Abstract

The ability to reliably discriminate vocal expressions of emotion is crucial to engage in successful social interactions. This process is arguably more crucial for blind individuals, since they cannot extract social information from faces and bodies, and therefore chiefly rely on voices to infer the emotional state of their interlocutors. Blind have demonstrated superior abilities in several aspects of auditory perception, but research on their ability to discriminate vocal features is still scarce and has provided unclear results. Here, we used a gating psychophysical paradigm to test whether early blind people would differ from individually matched sighted controls at the recognition of emotional expressions. Surprisingly, blind people showed lower performance than controls in discriminating specific vocal emotions. We presented segments of non-linguistic emotional vocalizations of increasing duration (100 to 400ms), portraying five basic emotions (fear, happy, sad, disgust, angry), and we asked our participants for an explicit emotion categorization task. We then calculated sensitivity indices and confusion patterns of their performance. We observed better performance of the sighted group in the discrimination of angry and fearful expression, with no between-group differences for other emotions. This result supports the view that vision plays a calibrating role for specific threat-related emotions specifically.

## Introduction

Achieving successful interactions with our conspecifics is a crucial skill to possess for humans and other social animals (Darwin, 1872). Such success is notably mediated by the efficient interpretation of a variety of sensory signals. In particular, facial, bodily and vocal signals convey a wealth of information about our interlocutors, with such information often being redundant across the senses. For example, identity, affect, age, gender, trustworthiness or speech can all be inferred through faces as well as through voices (Lange et al., 2022; Perrodin et al., 2015; Young et al. 2020). Ample evidence exists that neurotypical individuals rely more heavily on one sense compared to the others when performing specific inference on social information, with vision being the sense that is considered the most reliable for a large variety of perceptual tasks like the discrimination of age, emotion and identity (Amilon et al., 2007; Collignon et al., 2008; Hanley et al., 1998).

Despite its inherent multisensory nature, the perception of emotion expression is indeed known to be visually dominant. Research has shown that when differentiating between different basic emotions, visual cues override auditory cues across a wide range of emotions. Moreover, during incongruent audio-visual stimulation, observers tend to rely more on the emotions displayed through facial expressions rather than the voice when making decisions (Collignon et al., 2008). Additionally, when both facial and vocal expressions are presented simultaneously, participants’ errors are predominantly influenced by visual errors rather than auditory errors (Falagiarda & Collignon, 2019). These findings highlight the increased reliance on vision, even in situations where multiple senses are expected to provide optimal information. This stronger reliance on vision rather than audition is thought to relate to the enhanced salience of emotional features delivered through faces rather than voices (Collignon et al., 2008; Falagiarda & Collignon, 2019).

When vision has not been available throughout someone’s life, as it is the case for congenitally blind individuals, the reliance on vocal signals for most socially relevant information is arguably higher. Research on the perceptual consequence of early visual deprivation on the remaining senses has delivered multifaceted outcomes. Some research has shown that, because of the sensory deprivation, blind people perform better than sighted at a variety of non-visual tasks, such as sound pitch discrimination (Gougoux et al., 2004), sound localization (Battal et al., 2020) and fast speech discrimination (Moos & Trouvain, 2007). Such better performance in sensory deprived populations, at tasks carried out in their non-deprived senses, are often explained through sensory compensation, the idea that a sensory deprived individual must compensate for the lack of input in one sense by enhancing the computational capabilities of the remaining senses (Bavelier et al., 2006; Collignon et al., 2009). In contrast, deprived people have been shown to exhibit impairments in other perceptual tasks. For example, blind individuals show worse performance compared to sighted in auditory bisection (e.g. Gori et al., 2014). Such impairment is sometimes explained within the frame of sensory calibration, or rather lack thereof (Tonelli et al., 2015). A sensory calibration hypothesis posits that the dominant modality in a specific cognitive or sensory task is necessary during development to calibrate the other senses for that task. As a consequence, someone who lacked sensory calibration would perform more poorly, even when said task is performed unimodally in the non-dominant modality.

Interestingly, the sensory compensation and the sensory calibration hypotheses produce opposite predictions when looking at vocal discrimination of emotion in blind participants. Since emotion discrimination seems to be visually dominant, the lack of visual calibration might impair the optimal development of the categorization of vocal emotion expressions. On the other hand, lack of vision might be compensated by higher efficiency at extracting vocal information for successful social interactions, thus resulting in better performance. Research that has evaluated performance of the blind population on voice perception in general, and vocal expression of emotion in particular, remains limited and have produced conflicting results. For example, early blind people show better voice recognition abilities and learn new voice-name associations faster than sighted individuals (Bull et al., 1983; Hölig et al., 2014), while no performance difference was found between blind and sighted when asked to evaluate the body size of a speaker from their voice (Pisanski et al., 2016).

When it comes to emotional signals, developmental studies have shown that blind children and adolescents are worse and slower than the sighted when discriminating prosodies of emotional sentences (Chen et al., 2021) and have difficulties at recognizing vocal expressions of emotions (Blau, 1964; Dyck et al., 2004; Minter et al., 1991). Furthermore, blind children face challenges in emotion regulation (Chennaz et al. 2022) and have difficulties expressing emotions and reading other people’s mind (Roch-Levecq, 2006). In adults, Gamond et al. (2017) have tested the ability to detect a target emotion in vocalizations through a dichotic listening task, and found no accuracy difference between blind and sighted, while blind people were overall slower to respond. Martin and colleagues (2019) showed that participants with typical vision showed a greater advantage compared to blind participants when the emotional prosody used in a sentence matched its semantic content. In contrast, Klinge et al. (2010) found that the blind people were instead faster at responding to negative emotions in a discrimination task performed during fMRI scanning. The different results are likely due to differences in specific tasks employed, which might have not been appropriate to unveil group differences, or to the insufficient sensitivity of the experimental designs, which led to performance close to ceiling. Overall, these results depict an unclear picture, thus leaving the issue of better or worse performance of blind individuals in discrimination of voices, unresolved.

In the present study, we tested the auditory discrimination of a group of early blind individuals and a group of individually matched sighted individuals while categorizing vocal expression of a wide range of emotions. We considered our previously employed gating paradigm (Falagiarda & Collignon, 2019), an optimal way to test group differences, since the segmentation of the stimuli in different lengths creates a challenging task that has a higher chance of revealing subtle intergroup effects (Creupelandt et al., 2020). This paradigm assumes that when a perceiver performs the extraction of emotional information from voices, discrete stored properties of each emotion matching the incoming expression are rapidly and partially activated, similarly to the activation of the “word initial cohort” when hearing incoming speech (Marslen-Wilson, 1987). Our goal was to test whether early blind differs in the way they accumulate informational evidence at different time points during the unfolding of vocal emotional signals. In addition, we investigated whether discrimination follows a similar confusion pattern across our emotional set between the two groups.

## Methods

### Participants

The current study includes 16 early blind (EB) individuals and 16 sighted controls (SC) matched in age (±5 years) and sex (8 male subjects per group, mean age EB group = 34.7 years, mean age SC group = 34.1, no significant difference in groups’ ages: t = 0.19, p = 0.85), selected via opportunity sampling. Table 1 shows the information regarding the cause and age onset of blindness for the blind group. All participants had self-reported normal audition. The experimental protocol was approved by the institutional review boards of the University of Trento, Italy, and the Center for Mind/Brain Sciences (CIMeC). All participants provided a written informed consent prior to the experiment and received a monetary reimbursement for their participation.

**Table 1.**
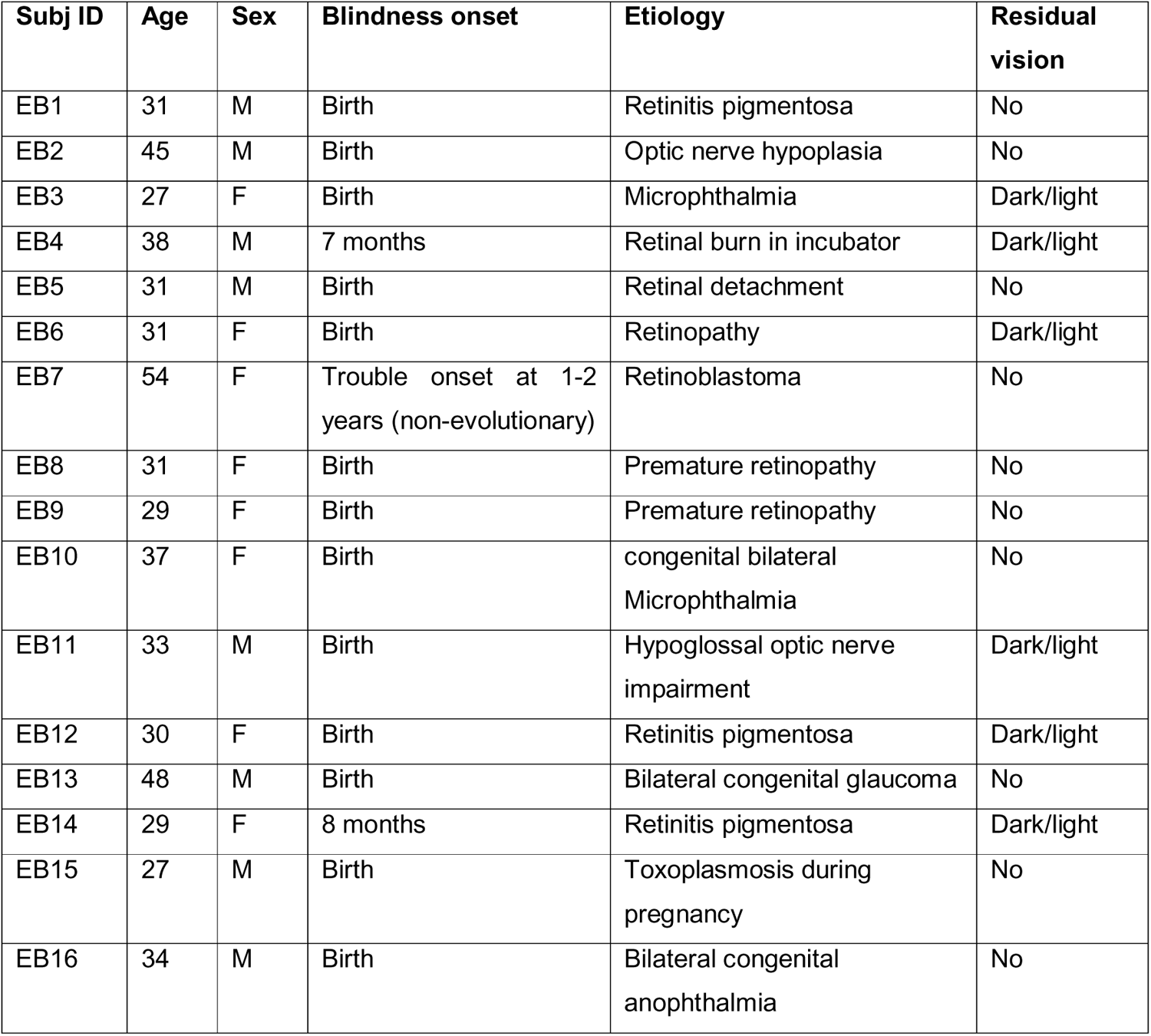
Etiology and blindness onset age of the blind group. Age at the time of testing and any residual light perception are also reported.

| Subj ID | Age | Sex | Blindness onset | Etiology | Residual vision |
| --- | --- | --- | --- | --- | --- |
| EB1 | 31 | M | Birth | Retinitis pigmentosa | No |
| EB2 | 45 | M | Birth | Optic nerve hypoplasia | No |
| EB3 | 27 | F | Birth | Microphthalmia | Dark/light |
| EB4 | 38 | M | 7 months | Retinal burn in incubator | Dark/light |
| EB5 | 31 | M | Birth | Retinal detachment | No |
| EB6 | 31 | F | Birth | Retinopathy | Dark/light |
| EB7 | 54 | F | Trouble onset at 1-2 years (non-evolutionary) | Retinoblastoma | No |
| EB8 | 31 | F | Birth | Premature retinopathy | No |
| EB9 | 29 | F | Birth | Premature retinopathy | No |
| EB10 | 37 | F | Birth | congenital bilateral Microphthalmia | No |
| EB11 | 33 | M | Birth | Hypoglossal optic nerve impairment | Dark/light |
| EB12 | 30 | F | Birth | Retinitis pigmentosa | Dark/light |
| EB13 | 48 | M | Birth | Bilateral congenital glaucoma | No |
| EB14 | 29 | F | 8 months | Retinitis pigmentosa | Dark/light |
| EB15 | 27 | M | Birth | Toxoplasmosis during pregnancy | No |
| EB16 | 34 | M | Birth | Bilateral congenital anophthalmia | No |

### Stimuli and design

The employed stimuli (available at http://osf.io/yedsz) consist of a set of non-linguistic vocal emotional expressions (i.e., vocalizations) representing the five basic emotions of anger, disgust, fear, joy and sadness, uttered by four different individuals (2 males). These exact stimuli have been used as auditory stimulation in our previous multisensory study conducted on the neurotypical population (see Falagiarda & Collignon, 2019; for a more detailed explanation of the stimuli selection process). We used a behavioral *gating* task (Grosjean, 1980) to present acoustic segments of incremental length (i.e., gate). The shortest stimulation segment is constituted by a silent audio clip lasting 100 ms. Participants were not informed of the presence of stimuli with no useful information for the purpose of the task, therefore they were asked to always provide an answer, even in the case of very short clips. Progressively longer gates were built through an increment of ∼33.37 ms to reach a maximum duration of stimulus presentation of 400 ms (i.e., gate 1: 100ms, gate 2: 133ms, gate 3: 167ms, … gate 10: 400ms).

### Procedure

Participants sat in a dimly lit room, in front of a laptop. Participants were additionally blindfolded prior to entering the room. The stimulation was presented through over-ear headphones (Hiearcool L1) at a comfortable, self-adjusted volume level. The task required from the participants was to identify the expressed emotion by pressing one of five buttons, where each button would correspond to one of the five possible emotion expressions (i.e., “anger”, “disgust”, “fear”, “joy”, “sadness”). At the beginning of the session, the experimenter would place the hands of the participants on the buttons required for the task, and play a pre-recorded file delivering the instructions. All participants would undergo a pre-experimental block aimed at ensuring that they had learned the mapping between each button and its assigned response. They also underwent a short practice block to familiarize with the experimental task.

During both pre-practice and practice blocks, participants received auditory feedback on the correctness of their answer, while no further feedback was given for the rest of the experiment. Each trial started with an interstimulus interval (ISI) that varied between 500 and 1000ms, followed by an auditory stimulus, selected randomly. After each stimulus, a tone was presented to signal to the participants that the 4s response time window had started; this was necessary to receive answers in all conditions, including those where no sound had been detected. Each emotion expression was portrayed by four different actors, therefore four trials in each emotion and each gate constituted a full repetition of the stimuli. Each participant was presented with two repetitions of the stimuli, for a total of 400 trials, presented as fully randomized in the two blocks of the experimental session. The experiment was run with Matlab 2013 using the Psychophysics toolbox extension (PTB-3, Kleiner et al., 2007).

### Transparency and openness

In compliance with APA endorsed Transparency and Openness Promotion Guidelines, we have reported above how we determined our sample size and data exclusions, and described all manipulations and measures we used in the study. We have shared the links to the stimuli, the data, analysis code, and research materials. Supplemental materials are available online and accessible via the journal website. Data were collected in 2017-18 and analyzed using R, version 4.2.1 (R Core Team, 2022); the packages used have been specified within the text. This study’s design and its analysis were not preregistered.

### Analyses and results

#### Sensitivity indices

We first discarded the trials in which the participants did not respond due to time out, or accidentally pressed a different button than the ones assigned for response (1.6% of the total). The rest of the data were used towards the computation of sensitivity indices (or *d’*, d-primes). The sensitivity indices constitute an unbiased measure of performance since they not only consider correct responses, but incorrect responses too (Signal Detection Theory, Macmillan & Creelman, 2004; Tanner & Swets, 1954). They are calculated as *d’ = z(Hit rate) – z(False alarm rate)*. The d’ were calculated for each subject, in each emotion expression at each gate (see Figure 1) and were subsequently analyzed to address our experimental question.

**Figure 1.**
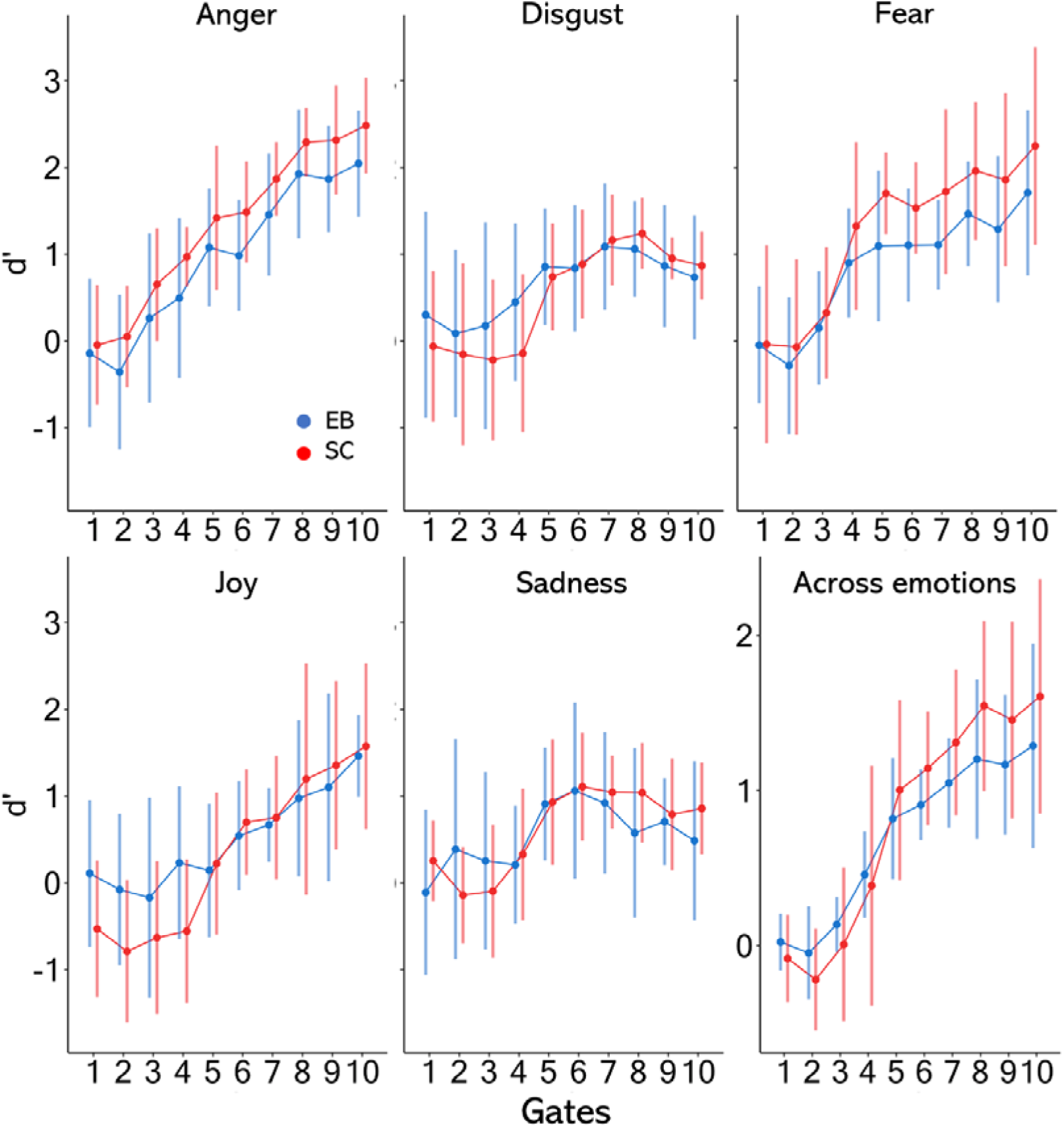
Sensitivity indices are here represented by gates and separately by group for each investigated emotion expression, as well as across emotions.

As previously mentioned, the shortest stimuli (gate 1) contained no emotional information, therefore the correspondent d’ values are theoretically meaningless and were removed from the following analyses. Outliers among the sensitivity indices were defined as the d’ values that exceeded 2.5 standard deviations (SD) from the mean of their condition, where one condition is defined by one of the five emotional categories in one gate for each group separately (cf. Falagiarda & Collignon, 2019). Nineteen values, corresponding to 1.3% of the d’ values, were found to be outlying and subsequently discarded. The remaining sensitivity indices were submitted to a generalized linear mixed model (GLMM) via the “lmerTest” package (Kuznetsova et al., 2017), with the predictors Group, Emotion and Gates as fixed effects, and Subject as random effect, and with the formula: *d-prime ∼ group * emotion * gate +(1|subj)* (marginal R^2^ = .46; conditional R^2^ = .51). In order to evaluate the global effects of the predictors, an analysis of variance was computed on the model through the “emmeans” package (Lenth, 2019) and using the Kenward-Rogers degrees of freedom approximation method. The ANOVA revealed a significant main effect of Emotion [F(4,1301.3) = 69.69, p < 0.001] and Gate [F(8, 1301.2) = 98.85, p < 0.001], as well as a significant interaction of Group by Emotion [F(4,1301.3) = 9.98, p < 0.001], Group by Gate [F(8,1301.2) = 2.97, p = 0.003], and Emotion by Gate [F(32,1301.1) = 5.13, p < 0.001]. The main effect of Group was non-significant [F(1, 30.0) = 2.1, p = 0.157], and so was the three-way interaction [F(32,1301) = 0.57, p = 0.97].

Post-hoc analyses on the main effect of Gate showed, as expected, a general increase in performance at the task as the stimulus duration increased, until reaching a plateau around gate 8, corresponding to a stimulus duration of 333ms. All tested gates were significantly different from each other, except gates 2 and 3; gate 5 and 6. Gate 7 was not significantly different from gate 6, 8 or 9; gate 8, 9 and 10 were not significantly different from each other (all significant p’s < 0.039; all other p’s > 0.24; all p’s adjusted through Tukey’s method for comparing a family of 9 estimates, see Figure 2A). Post-hoc analyses on the main effect of Emotion showed that anger and fear lead to significantly higher performance in the task than all other three emotions: anger vs disgust, anger vs joy, anger vs sadness, fear vs disgust, fear vs joy, fear vs sadness (p’s < 0.001). The d’ values for anger and fear were not significantly different from each other, and the same was shown for disgust, joy and sadness: anger vs fear, disgust vs joy, disgust vs joy, joy vs disgust (p’s > 0.083; all p’s adjusted through Tukey’s method for comparing a family of 5 estimates, see Figure 2B).

**Figure 2.**
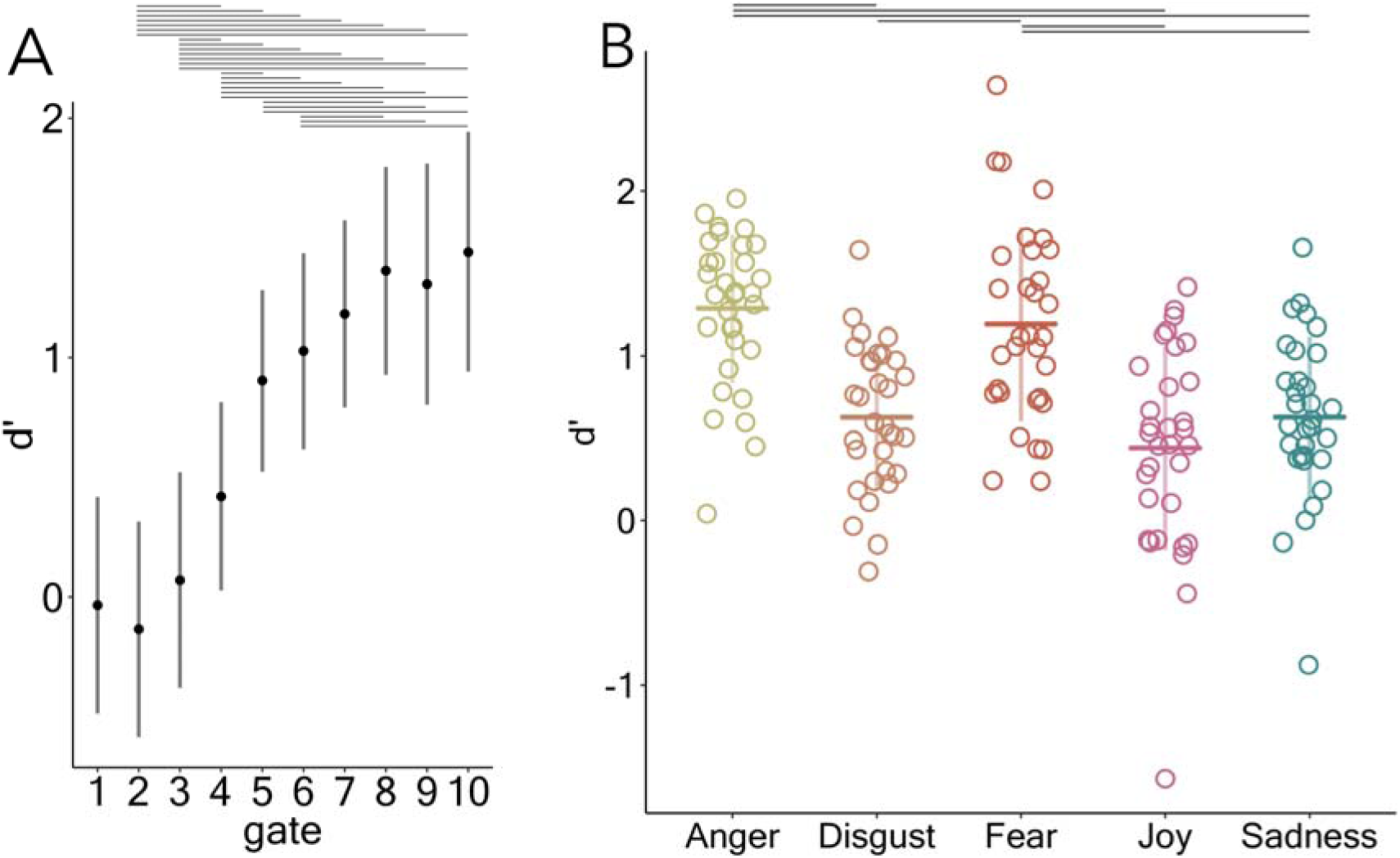
Representation of the main effect of gate [A] and the main effect of emotion [B]. In [A] sensitivity increases as the stimulus duration, i.e. the amount of information available, increases, reaching a plateau at gate 8 (333ms). In [B] each circle represents one participant; Anger and Fear lead to significantly higher performance than the other three emotion expressions. Error bars are the SDs.

Post-hoc analyses on the interaction between Group and Emotion aimed at five comparisons of interest, namely the comparison between the two groups within the same emotion category. Those comparisons revealed to be significant for anger and fear, where the sighted participants’ group exhibited overall higher sensitivity, hence higher performance than the blind group (anger: EB vs SC, p = 0.038, fear: EB vs SC, p = 0.015), but not for the other emotions (disgust, joy and sadness: EB vs SC, p’s > 0.99; all p’s adjusted through Tukey’s method for comparing a family of 10 estimates; see Figure 3.).

**Figure 3.**
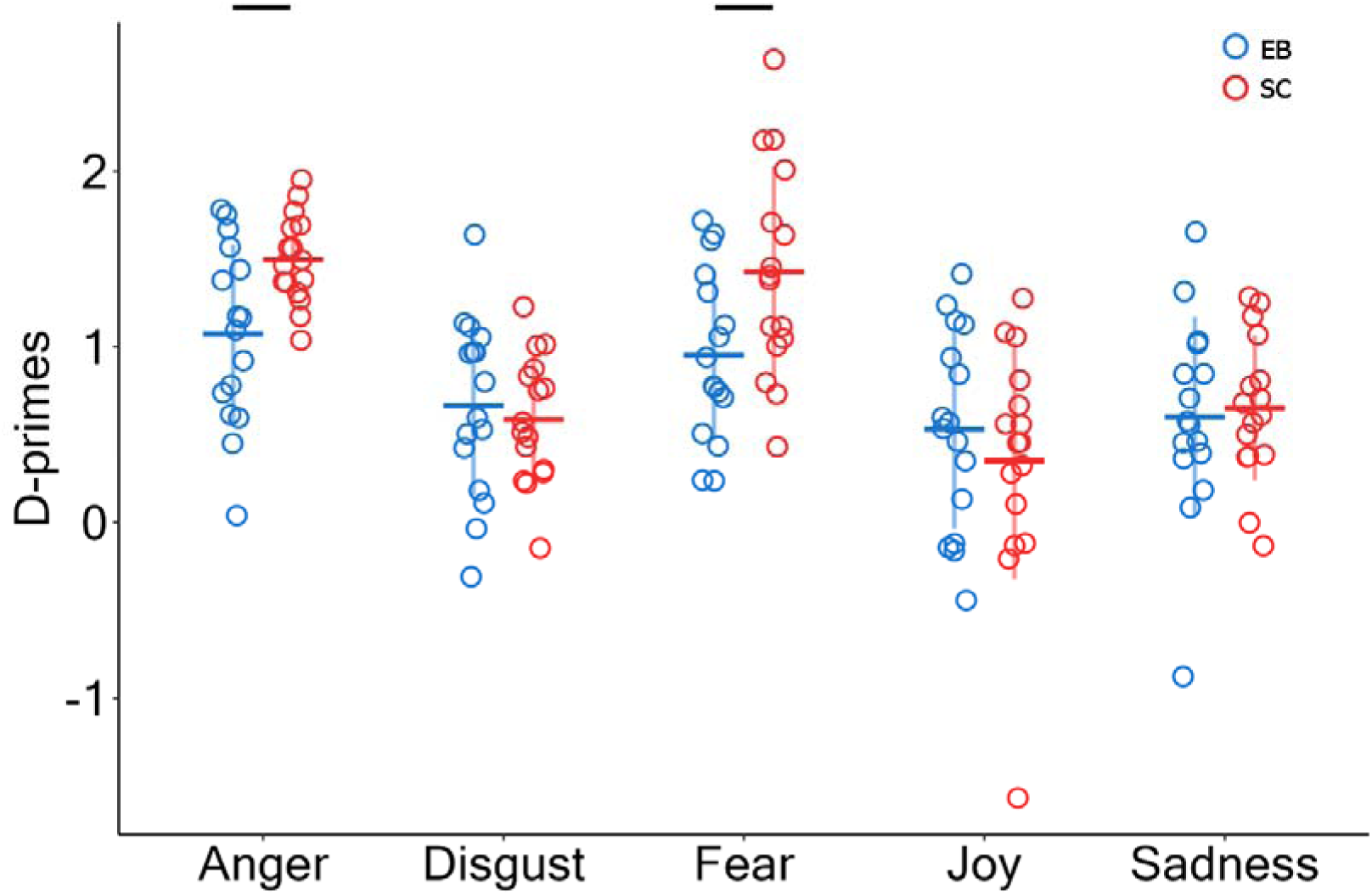
Representation of the interaction between emotion and group. In the expressions of anger and fear, sighted individuals exhibit significantly higher performance compared to blind subjects. Each circle represents one participant. Error bars are the SDs.

Post-hoc analyses on the interaction between Gate and Group failed to highlight any significant difference in the performance of the two groups across the gates, all p’s > .05 (see Figure 1). Lastly, the interaction between Gate and Emotion pointed to a different increase in performance at the increased stimulus length in the different emotions, which means that the amount of information at each gate differed across vocal expression of emotions. The thorough interpretation of each potentially significant post-hoc pairwise comparison for these two interactions is beyond the scope of this investigation and will not be discussed further.

#### Confusion matrices

To test whether the behavior of sighted and blind individuals was similar when producing incorrect responses, we made use of confusion matrices. Confusion matrices were calculated for each subject with the responses at each investigated time point separately. In confusion matrices, the diagonal represents the correct responses, while all the data laying off the diagonal represents incorrect responses, or confusion patterns (see Figure 2 for the color-coded average matrices of the two groups at each gate).

Through confusion matrices, we wanted to test whether the two groups behaved similarly when producing incorrect responses; in particular, we wanted to test whether their incorrect responses would be more similar within one group compared to the other group. A significant difference would hint at a partially different strategy between the groups when completing the task. To address this question, we calculated an intragroup and an intergroup measure of correlation at each gate. The intragroup correlation was computed by calculating Pearson’s correlation coefficient between the confusion patterns (on-diagonal data discarded; see Ritchie et al., 2017) of each subject with the average confusions of the rest of his/her own group (blind or sighted). The intergroup correlation was instead computed by correlating each subject to the mean of the other group. In order to reveal any difference, we tested the average intragroup vs the average intergroup correlation coefficients (Fisher transformed Pearson’s r). No significant difference was found at any gate (t-tests, all p’s_FDR_ > 0.39). Failing to find a significant difference between intragroup and intergroup correlations supports to the idea that the incorrect responses are similar in the two groups.

Furthermore, we tested the intergroup correlation against zero. The results showed significant coefficients for gate 1, and gate 4 to 10. Gate 1 and 4 showed a weak correlation coefficient (gate 1: r_FisherZ_ = 0.15, t = 2.83, p_FDR_ = 0.01; gate 4: r_FisherZ_ = 0.16, t = 3.43, p_FDR_ = 0.002), while gates 5 to 10 showed moderate to strong degrees of correlation (gate 5: r_FisherZ_ = 0.38, t = 6.4, p_FDR_ < 0.0001; gate 6: r_FisherZ_ = 0.4, t = 7.84, p_FDR_ < 0.0001; gate 7: r_FisherZ_ = 0.48, t = 11.16, p_FDR_ < 0.0001; gate 8 r_FisherZ_ = 0.58, t = 10.83, p_FDR_ < 0.0001; gate 9: r_FisherZ_ = 0.54, t = 12.13, p_FDR_ < 0.0001; gate 10: r_FisherZ_ = 0.61, t = 12.41, p_FDR_ < 0.0001). No significant correlation was found for gate 2 and 3 (gate2: r_FisherZ_ = 0.09, t = 1.62, p_FDR_ = 0.13; gate 3: r_FisherZ_ = 0.09, t = 1.34, p_FDR_ = 0.19). These results showed an increasingly higher similarity between the incorrect responses (the confusions) of the two groups for stimuli of 200ms and longer (gate 4 and above), as well as a small degree of similarity of the confusions for the shortest stimuli (gate 1). At gate 1, where no emotional information was actually presented, the correlation is probably given by a similar consistency in the responses that subjects adopted. Looking at the color-coded matrices in Figure 3, it seems that, under such high uncertainty context (no actual information is available to complete the task), there might be a potential bias to report disgust. From gate 4 onwards, when, arguably, enough information is delivered to lead to a more consistent emotion categorization process, the similarity in the confusion patterns raised, showing that blind and sighted confused vocal emotions at a similar extent. Based on a mere visual evaluation of the matrices, we might notice that joy seems to be misclassified as sadness in both groups, more pronouncedly around the middle of the signal (see Figure 3). All correlation values are summarized in Figure 4, and their distributions represented in Figure S1.

**Figure 4.**
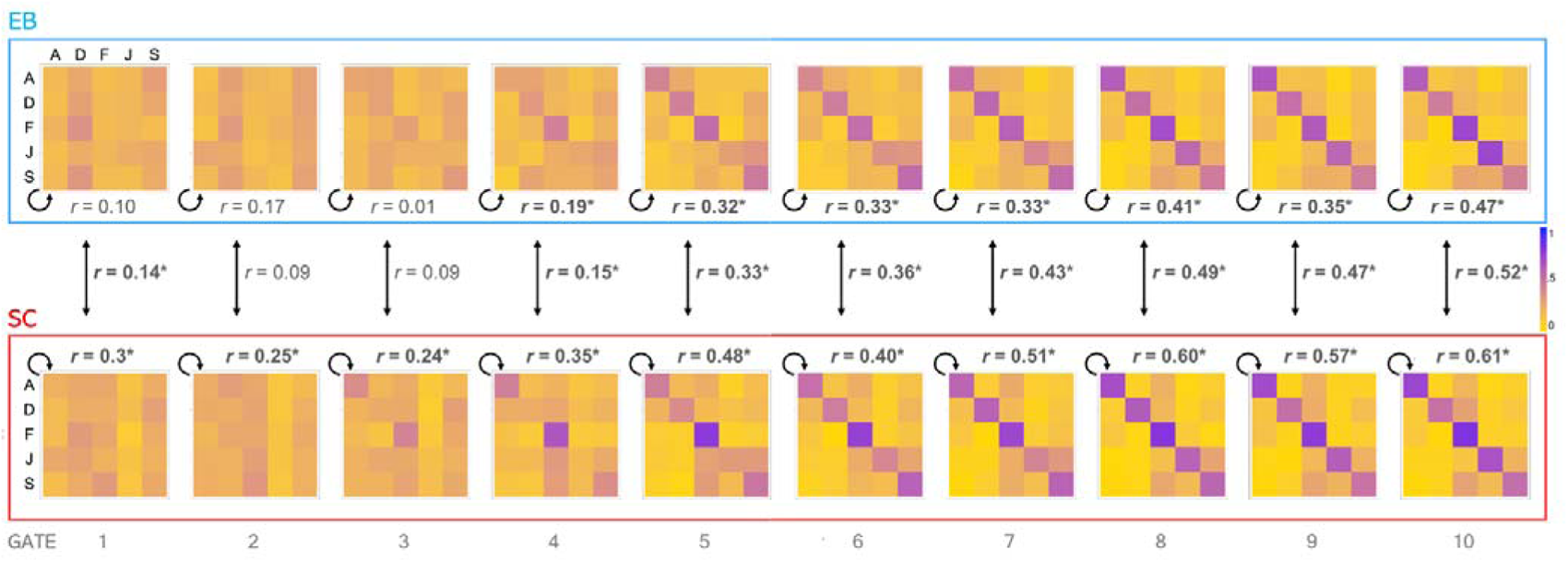
Average confusion matrices by group and by gate. The emotions contained in the stimuli are represented by rows, while the emotions of the subjects’ responses are represented by columns (A = anger, D = disgust, F = fear, J = joy, S = sadness). The diagonal of each matrix represents the correct responses, off-diagonal data represents the actual confusion patterns. Between the two matrices at each gate is the average inter-group correlation coefficient (here reported untransformed), while next to each matrix is the value of the intra-group correlation of each group. Significance is assessed on Fisher’s transformed coefficient values.

We also confirmed our results using a non-parametric, measure of sensitivity (A-prime; Zhang and Mueller, 2005) as supplemental material.

## Discussion

In the present study, we compared the ability of sighted and blind individuals at categorizing emotions, as perceived through vocal non-linguistic segments of different lengths. The gating paradigm we used relies on extracting emotional information from vocal expressions in conditions where information about emotions is progressively accumulated over time. The challenging condition imposed on the perceptual system avoids any ceiling effects and therefore allows for sensitive measurement of the abilities to efficiently extract emotionally relevant acoustic information. Our results show that sighted individuals outperformed the blind at discriminating anger and fear, while no difference between the groups was found for expressions of disgust, joy or sadness. Furthermore, we observed that, independently of the sensory experience of our participants – blind or sighted – the threat-related auditory expressions of anger and fear were recognized better compared to the other investigated categories (Figure 2B).

In past research, threat-related emotions, with a particular attention given to fear, have proved to stand out when compared to other basic emotional categories (Damasio & Carvalho, 2013). For example, facial (Milders et al., 2006) or vocal (Falagiarda & Collignon, 2019) expressions of fear are detected or discriminated better than other basic emotions. Also, fearful faces more easily and preferentially break through conscious perception, whether evaluated through binocular rivalry (Amting et al., 2010), or continuous flash suppressions (Yang et al., 2007). Four months old infants as well as adults exhibit an avoidant looking behavior when presented with angry and fearful faces, but not with happy, sad or neutral faces, highlighting how the recognition of threat is present very early in life, and that threat is an attribute allegedly independent from valence (since sadness, like anger and fear, is also negative in valence, but did not trigger avoidance; Hunnius et al., 2011). These conclusions can be extended to the domain of threat-related vocal signals, as newborns of just a few days of age show higher and differential mismatch responses for fearful and angry vocalizations compared to happy and neutral ones (Cheng et al. 2012). These results altogether highlight the importance of these threat-related emotions, that arguably resides in their link to fast responses towards survival, a “fight-or-flight” type of responses.

Our results suggest a potential calibrating role of visual experience for the optimal development of the perception of threat-related vocal expressions. Why is the discrimination of fear and anger vocal expressions specifically susceptible to calibration by visual experience? As described above, fear and anger expressions show the highest discrimination when compared to other emotions, meaning that they need less presentation time for a correct discrimination to be reached. Interestingly, this is also true for facial expressions of emotions (Falagiarda & Collignon, 2019). This might relate to the importance of quickly and efficiently recognize these two emotions for survival in humans and other animals, with evolution creating pressure for their visual expression being very salient (Damasio & Carvalho, 2013). One might therefore speculate that multisensory perceptual learning and calibration processes are particularly important for those emotion. Indeed, perceptual learning studies have shown that audiovisual training reflects in a more refined discriminating of unimodal auditory stimuli (Lidestam et al., 2014; Moradi et al., 2019). It might therefore be that the optimal development of the discrimination of anger and fear through voices more specifically depend on their coupling with facial expression through development (Young et al., 2020), something that is absent in early blind people. Throughout their lives, sighted people are massively exposed to sensory stimulation being simultaneously delivered via visual and auditory channels (Klasen et al., 2012). Being able to pool together facial and vocal cues and combining them with contextual information is of pivotal importance in emotion interpretation, due to the highly changeable and ambiguous nature of interpersonal interactions (Theurel et al., 2016; Young et al., 2020). Unlike the sighted, early blind people cannot rely on extensive training at integrating these different sources of information, resulting in a detrimental impact on their capability to interpret emotional states from vocal inputs (Minter et al., 1991; Valente et al., 2018). A further avenue to gather evidence on the effect of visual calibration could be the investigation of late onset blindness: late blind individuals have lost the sense of sight later in life, implying that vision was present at the critical period needed for calibration. A sensory calibration hypothesis would therefore not predict an impairment at a vocal discrimination task in late blind compared to early blind subjects (cf. Chen et al., 2021).

Another, not mutually exclusive, speculation to explain the specific impairment in the discrimination of anger and fear expressions in blind people may relate to a lower exposure to threat-related situations in blind people. Modern societies are almost never designed for the comfortable fruition of people with disabilities and impairment. We could speculate that blind individuals grew up in a more “protected” environment than their non-visually impaired peers. They have therefore been potentially less exposed to threatening situations, and the people around them might have been less inclined to express anger or fear in their presence. If this were true, they might have overall lower expertise with rageous and frightened vocal utterances, resulting in lower discriminative abilities. A further account worth considering here relates to the concept of embodiment of emotion, according to which perceiving and thinking about emotion automatically activate motoric re-experience of emotion in one’s self (Hari & Kujala, 2009; Kragel & LaBar, 2016; Niedenthal, 2007). For instance, the presentation of facial expressions of emotion triggers the motor recreation of the perceived facial expression in the observer’s own face, known as facial mimicry (Niedenthal et al., 2010). The embodied processing of facial expression contributes to the recognition and understanding of emotions, thus playing a role as social regulation (Beffarra et al., 2012; Hess & Fisher, 2013). Interestingly, embodied responses have been observed for auditory stimulation (Fino et al., 2016; Korb et al., 2015), including nonverbal emotion vocalizations (Hawk et al., 2012). One might argue whether similar effects could be observed in early blind, a population who have not been exposed to facial expressions of emotion. Could facial mimicry be observed in people lacking any modelling experience in how to express emotions via facial cues? Studies on this topic suggest that, although visual learning does not seem to be a prerequisite to exhibit patterns of facial expressions, some variations in the control and intensity of emotions have been noticed in the blind population (see Valente et al., 2018 for a review). In particular, blind individuals conform less to the display rules dictating what emotions and levels of intensity is socially acceptable to show in different social contexts (Galati et al., 2003; Kunz et al., 2012). Thus, the possibility holds that blind individuals, not having been exposed to visual expressions of emotion during their life, might be impaired in their capability to perform motor simulations of such expressions, mimic such expressions of emotion (although see Arias et al., 2021), which, in turn, can reflect in poorer vocal emotion discrimination in this population.

One might argue whether the different emotion socialization process blind and sighted individuals have experienced during their development might have resulted in the use of distinct strategies by the two groups while performing our task. To answer this question, we performed a closer inspection of their incorrect responses, or confusions, to highlight potential emerging patterns. We represented their errors as confusion matrices and computed intergroup correlations for each of the ten stimulus durations of the design. This analysis revealed increasingly higher degrees of correlation for stimuli starting at 200ms and above (gates 4-10; see Figure 4). This indicates that the two groups confuse the same emotions when responding incorrectly. The idea that similar discriminative strategies are employed by the two groups is supported also by the comparison of the intergroup correlations with the intragroup correlations, which did not show any differences at all gates, further supporting the idea that the incorrect performances of the two groups are highly aligned, and no group-specific sub-patterns of responses seem to exist. Based on these results, we speculate that blind and the sighted people engage similar perceptual processes to classify emotions. If the responses in uncertain circumstances, that later lead to incorrect discriminatory decisions, are driven by similar factors in the blind and sighted groups, we can argue that these factors should not include “visual” strategies (e.g. visual imagery), since the blind would not be able to implement them. We would therefore suggest, firstly, that a certain amount of information needs to unfold in time before perceptual elements get extracted in a consistent manner across individuals (in the present design, 200ms of stimulation), and secondly, that these perceptual elements used towards a discriminatory decision are acoustic features of the vocal emotional expressions. Whether the results we observed are specific to the emotional domain, or they rather reflect the way blind people base their decisions on accumulating auditory information in general, is beyond the scope of the present paper and could be matter of future investigation. Along the same line, in accordance with recent evidence showing that 24 distinct emotional states can be extracted from non-linguistic vocalizations (Cowen et al., 2019), it would be interesting to test more fine-grained emotional states in addition to the primary emotions tested here.

In conclusion, we investigated the abilities of early blind and sighted individuals at a vocal emotion discrimination task based on stimuli of incremental length. An overall higher sensitivity for expressions of anger and fear was found when compared to other emotions. Additionally, sighted participants performed better than blind at discriminating those same two threat-related emotional categories, while no difference was found for expressions of disgust, joy or sadness. These results show firstly how angry and fearful expressions set themselves aside from other basic emotions, extending results in vision (Marsh et al., 2005a, 2005b; Pichon et al., 2009; Sacco & Hugenberg, 2009) in the less investigated auditory domain (but also see Erlich et al., 2013; Scott et al., 1997). Secondly, these results suggest that facial inputs are required for the fine tuning of discrimination of threat-related expressions. Lastly, we observed similar pattern of confusions across emotions in the two groups once enough signal has unfolded over time, suggesting the adoption of similar and allegedly non-visually based strategies while performing the task. Even if speculative, we suggested mechanistic explanations as to why the discrimination of vocalization of fear and anger suffers from a lack of visual experience.

## CRediT author statement

Federica Falagiarda: Conceptualization, Data curation, Methodology, Software, Formal Analysis, Investigation, Writing– Original Draft, Visualization

Olivier Collignon: Conceptualization, Methodology, Writing-Review and Editing, Supervision, Project Administration, Funding Acquisition

Valeria Occelli: Investigation, Data curation, Writing – Review & Editing, Funding acquisition

## Acknowledgements

This work was supported by a European Research Council starting grant (MADVIS grant #337573) attributed to Olivier Collignon and by a European Union’s Horizon 2020 research and innovation programme under the Marie Sklodowska-Curie grant agreement (#701250) awarded to Valeria Occelli. Federica Falagiarda is a research fellow and Olivier Collignon a research associate at the National Fund for Scientific Research of Belgium (FRS-FNRS).

## Openly available data

The data supporting the findings reported in this paper as well as the scripts used in the analyses are openly available from the Open Science Framework repository at the link: https://osf.io/4zksp/.

