## Supplemental Material for "Vision plays a calibrating role in discriminating threat-related vocal emotions"

Supplementary material

Methods

The current gating paradigm (see main Methods section), was originally designed for the psychometric fitting of performance-based curves. Such analyses and the reasons why they are not reported in the main paper are explained in the following sections.

Supplemental results

Psychometric curve fitting

We decided to move away from psychometric fitting and instead rely on the calculation of d’ for the following observations. Firstly, by the outliers’ definition followed in our first publication with such paradigm (see Falagiarda and Collignon, 2019), the exclusion rate surpassed 13% of the calculated thresholds (see Table S1).

Secondly, the results yielded by the analyses, conducted after outliers’ exclusion, was difficult to interpret towards either one conclusion or the opposite when it came to group comparisons. You can find such results reported in the next section.

Table S1. Thresholds outliers defined through goodness of fit testing, visual inspection of the curves, and values exceeding 2.5SD from the mean of their condition.

Subject numbers between 1 and 16 refer to blind subjects, between 17 and 32 to sighted controls.

| **Emotion** | **Bad fit/decreasing curve (visual inspection)** | **+/- 2.5SD from mean of condition** | **Total count** |
| --- | --- | --- | --- |
| Anger | Subj nr: 4 | 21 | 2s |
| Disgust | 1, 13, 14, 23 | 16, 32 | 6s |
| Fear | 1, 27 |  | 2s |
| Joy | 5, 27 | 11 | 3s |
| Sadness | 1, 12, 15, 16, 23, 24 | 9, 17 | 8s |
| Overall count | 15s | 6s | 21s |

Thresholds were obtained through the fitting of logistic curves to the data of each participant for each emotion condition separately (goodness of fit evaluated through a bootstrap procedure, n = 200), for a total of 160 fitting procedures.

A generalized linear mixed model (GLMM) was run on the 139 values that were not identified as outliers (see outliers’ definition in table S1), with the predictors Group and Emotion as fixed effects, and Subject as random effect, and with the formula: *threshold ~ group * emotion +(1|subj)*. In order to evaluate the global effects of the predictors, an analysis of variance was computed on the model (Kenward-Rogers degrees of freedom approximation method). The ANOVA revealed a trend in the main effect of group (F = 4.11, p = 0.052), the main effect of emotion was significant (F = 12.75, p < 0.0001) while the interaction between emotion and group was non-significant (F = 1.98, p > 0.1).

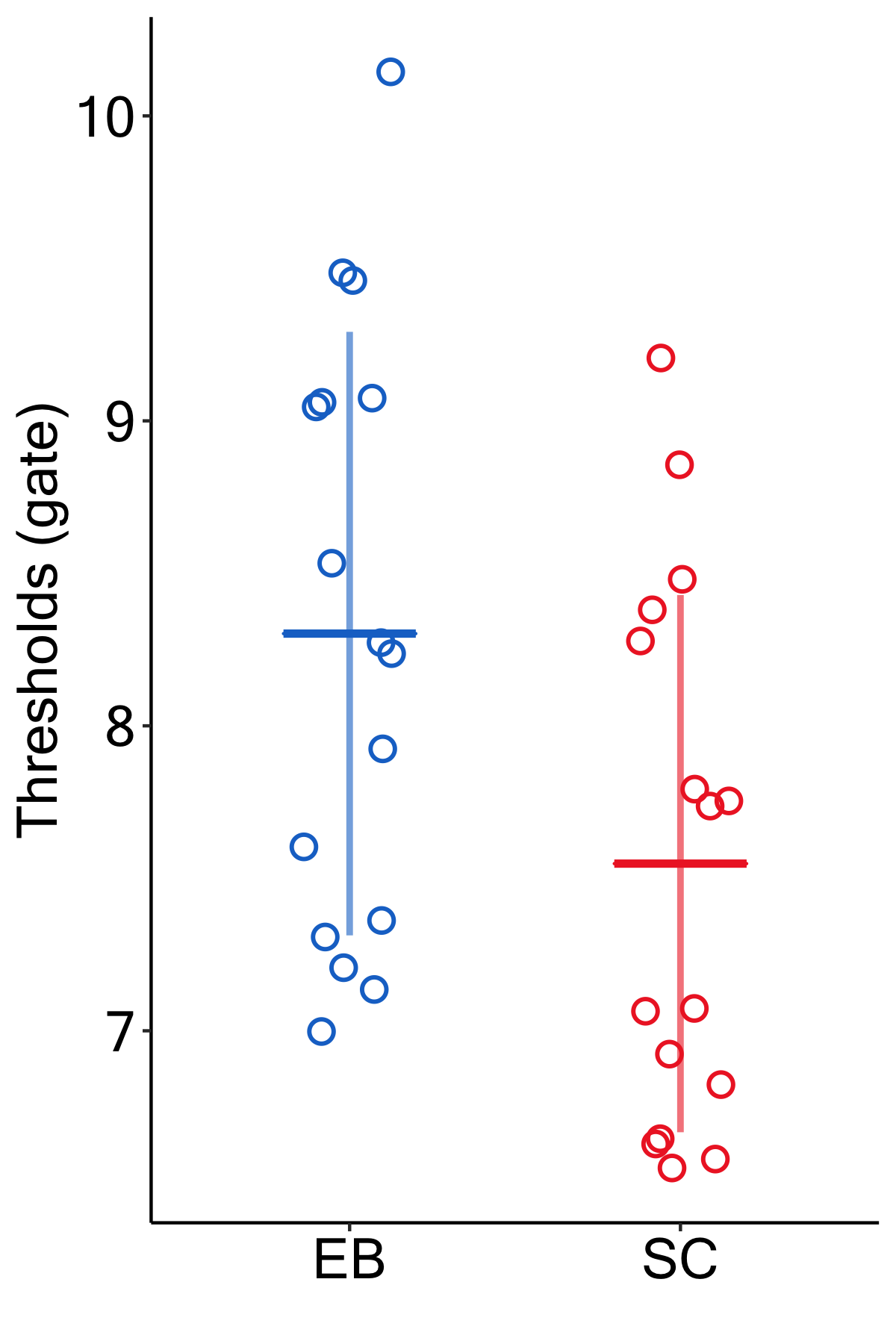

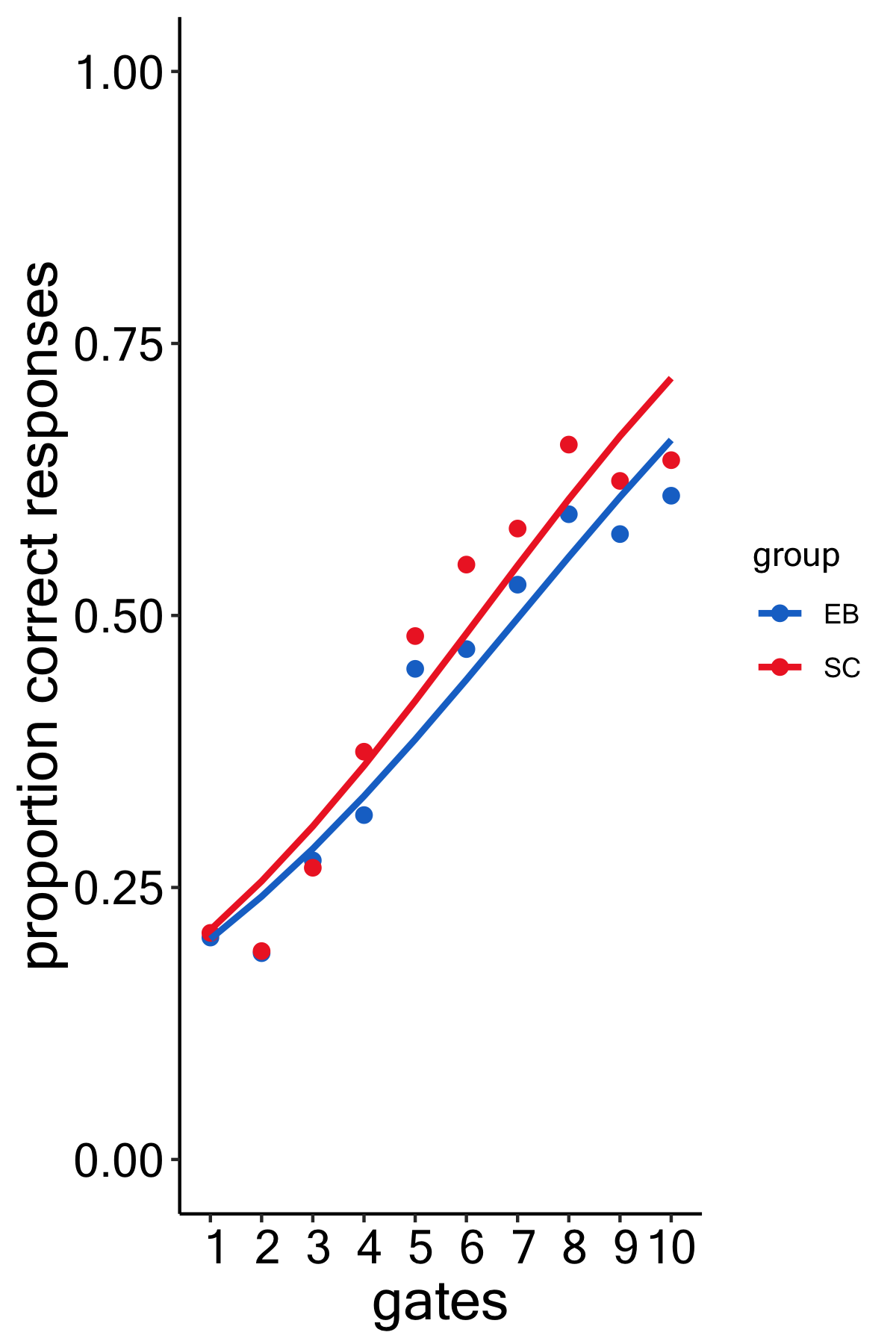

Figure S1. The thresholds’ distributions (left) in the two groups, and fitted curves (right) across emotions, data of the entire group pooled together for visualization purposes only.

Confusion matrices correlations’ distributions

The confusion matrices (see main figure 5) were correlated in the following fashion: after excluding the on-diagonal data, the matrix of each subject was correlated to the average of the rest of that subject’s group in order to obtain an intra-group correlation value, and then to the average of the opposite group in order to obtain an inter-group correlation value (see main paper for the results of this analysis). The distributions of the said correlations are represented here in figure S2.

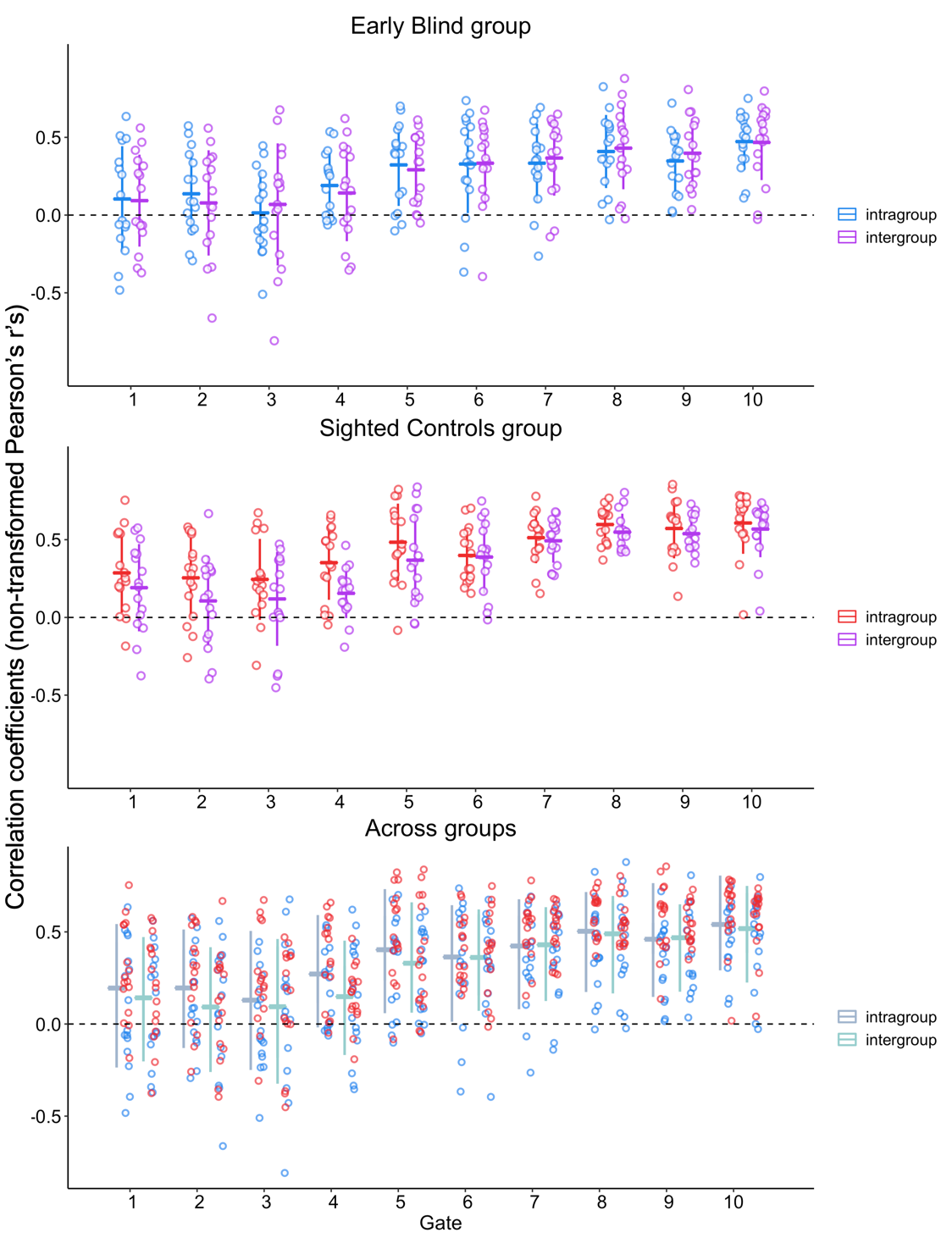

Figure S2. The distributions of the untransformed intra- and inter-group correlations. The coefficients are represented separately by group on the top two panels, while pooled together in the bottom panel. The color of the dot in the third panel refers to the subject whose individual data was used for the correlation (whether intra- or inter-group), with blue referring to a blind participant and red to a sighted control.

Average confusion matrices correlations

As a further measure of (inter-group) correlation, we tested testing the group-averaged confusion matrices. This leads to strong and significant correlation coefficients for gate 5 and above (gate 5: r = 0.652, p = 0.003; gate 6: r = 0.765, p = 0.0002, gate 7: r = 0.849, p < 0.0001; gate 8: r = 0.853, p < 0.0001; gate 9: r = 0.856, p < 0.0001; gate 10: r = 0.847, p = ; gates 1-4 all p’s > 0.117; all reported p’s are FDR-corrected).

A-prime scores

In addition to d-prime scores, we calculated an alternative, non-parametric, measure of sensitivity, A-prime scores (Zhang & Mueller, 2005). Twenty-two values, corresponding to 1.5% of the values, were found to be outlying and subsequently discarded. The remaining sensitivity indices were submitted to a generalized linear mixed model (GLMM), with the predictors Group, Emotion and Gates as fixed effects, and Subject as random effect, and with the formula: *A-prime ~ group * emotion * gate +(1|subj*. The ANOVA conducted on the effects of the predictors revealed a significant main effect of Emotion [F(4, 1298.2) = 69.50, p < 0.001] and Gate [F(8, 1298.2) = 105.90, p < 0.001], as well as a significant interaction of Group by Emotion [F(4, 1298.2) = 6.11, p < 0.001], Group by Gate [F(8, 1298.2) = 2.98, p = 0.003], and Emotion by Gate [F(32,1298.1) = 4.51, p < 0.001]. The main effect of Group was non-significant [F(1, 30.0) = 1.58, p = 0.219], and so was the three-way interaction [F(32, 1298.1) = 1.03, p = 0.42]. A-prime scores are represented by gates and separately by group for each investigated emotion, as well as across emotions, in Figure S3.

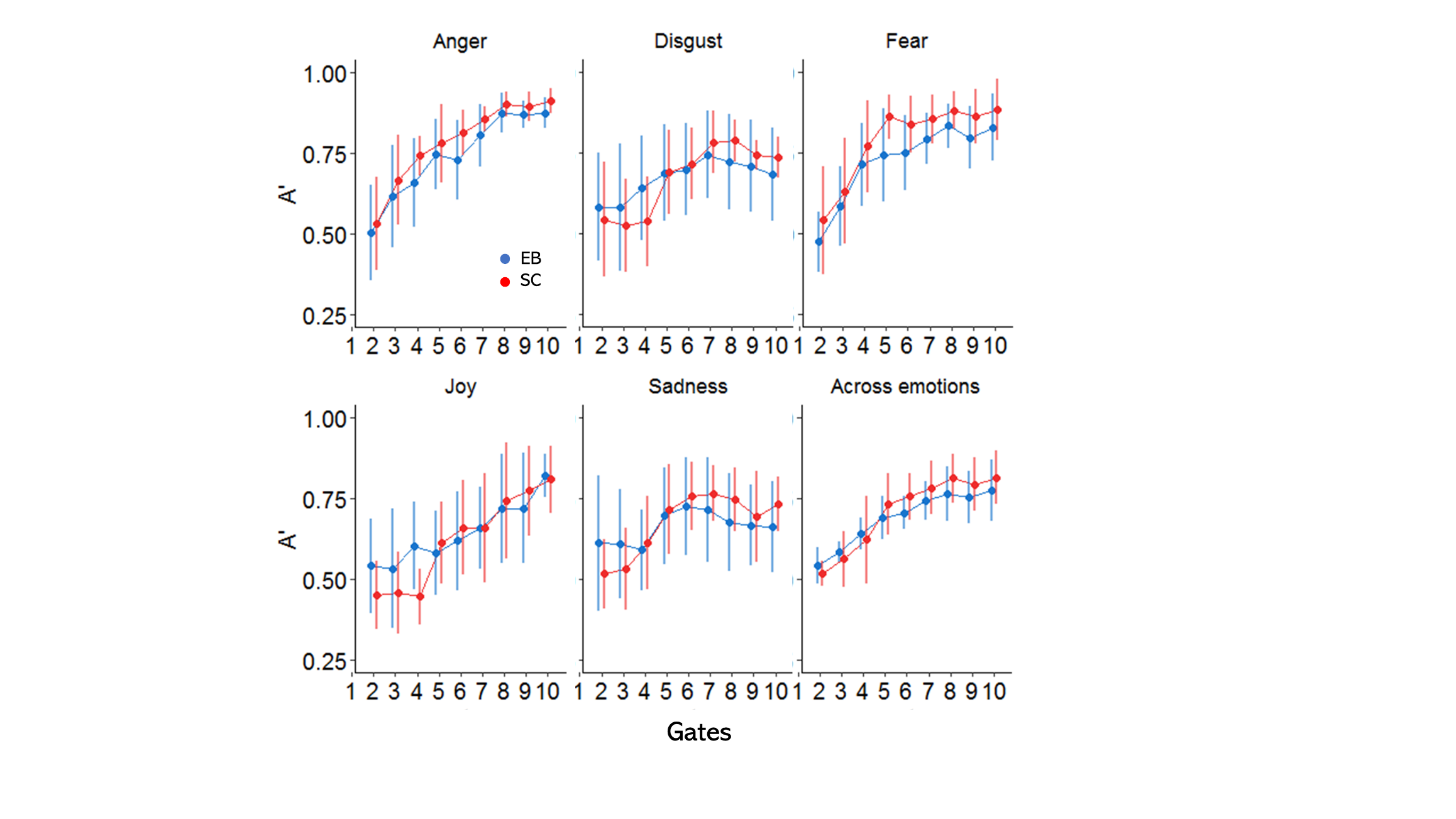

Figure S3. A-prime scores are here represented by gates and separately by group for each investigated emotion expression, as well as across emotions.

Discussion

The results of the analysis of the psychometric thresholds of discrimination were inconclusive in regard of a group difference between blind and sighted when perceiving short bursts of vocal emotions. The trend seemed to suggested a better performance (lower thresholds) of the sighted group (see fig S1), however we did not feel confident in making such a claim with only a trend in the statistical results. Neither did we feel confident in making the opposite claim – of a non-detected difference – based on that very same result.

The analysis of a-prime scores mostly mirrored that obtained by using d-prime scores described in the main text.
